# tbx5 and tbx15 mediate zebrafish posterior lateral line migration and development via a negative feedback loop

**DOI:** 10.1101/452797

**Authors:** Heather R. Whitehurst, Viviana E. Gallardo, Shawn Burgess

## Abstract

The zebrafish posterior lateral line (pLL) is a mechanosensory organ enabling detection of movement in the aqueous environment. Evidence suggests that two T-box transcription factors, *tbx5* and *tbx15*, participate in pLL migration and development. Lateral line migration and deposition defects are observed following *tbx5* or *tbx15* morpholino knockdown. Additional studies demonstrate that the *tbx5* phenotype is partially rescued when *tbx15* is also knocked down. pLL defects similar to those of *tbx5* are noted following knockdown of *camKII-ß2*, a known downstream target of Tbx5, suggesting a potential mechanistic pathway. Ectopic expression of human *CamKII-∂C* partially rescues the *tbx5* phenotype but exacerbates the *tbx15* phenotype. These results, combined with in situ hybridization and qRT-PCR profiling, indicate a negative feedback loop controlling multiple primordium patterning markers, including *cxcr4b, cxcr7b, fgf10a* and *notch3*, which ultimately affects pLL neuromast migration and deposition.

## Introduction

The zebrafish (Danio rerio) lateral line system is subdivided into the anterior (aLL) and posterior lateral lines (pLL). At 3 days post-fertilization (dpf), the pLL is comprised of 7-9 mechanosensory patches, called neuromasts, spaced at regular intervals along the horizontal myoseptum (Raible and Kruse, 2000; Ghysen and Dambly-Chaudiere, 2007). During the initial formation process, beginning at about 24 hours post fertilization (hpf), the pLL primordium forms just posterior to the otic placode, organizes into a leading zone and a trailing zone (Metcalfe et al., 1985), and migrates anterior to posterior along the myoseptum, depositing neuromasts periodically (Gompel et al., 2001; Haas and Gilmour, 2006). The leading zone includes progenitor cells and cells undergoing maturation to become rosettes (Laguerre et al., 2005; Laguerre et al., 2009). The trailing zone contains rosette structures, which are neuromast precursors (Nechiporuk and Raible, 2008). Maturation and deposition occur in a stepwise manner where the most mature rosette is deposited as new cells destined to form additional rosettes are added to the primordium by progenitor cells in the leading zone (Nechiporuk and Raible, 2008). The primordium deposits 5-6 primary neuromasts along the trunk of the fish then continues its journey to the tail, fragmenting into an additional 2-3 neuromasts (Gompel et al., 2001; Harris et al., 2003; Haas and Gilmour, 2006).

The two genes of interest for these studies, *tbx5* and *tbx15*, are both traditional t-box transcription factors. This family of transcription factors is recognized for roles in development and is characterized by a DNA binding domain consisting of a core T-site (Conlon et al., 2001; Takeuchi et al., 2003a). Specificity is partially conferred by the arrangement of the t-site in tandem or inverted repeats and the number of core T-box sequences that compose the binding site. Additional specificity is generated by diversity in binding partners. T-box transcription factors can function as monomers, homodimers or heterodimers, thus allowing for multiple functions for a single Tbx gene (Conlon et al., 2001; Kiefer, 2004).

Tbx5 is associated with human Holt-Oram syndrome, characterized by skeletal and cardiac defects that are widely variable in severity but generally affect the forelimbs, usually to a greater extent on the left side (Li et al., 1997). In zebrafish, *tbx5* has roles in development of both the heart and the pectoral fins (Ruvinsky et al., 2000; Garrity et al., 2002; Takeuchi et al., 2003a; Takeuchi et al., 2003b). *Heartstrings* mutants lack pectoral fins and their heart fails to properly loop and form an atrium and ventricle, leading to edema in the developing heart chambers and eventually death (Garrity et al., 2002).

*Tbx15* is the gene involved in Cousin syndrome in humans, which is characterized by short stature, craniofacial dysmorphisms, including low-set and rotated ears with hypoacusia, and hypoplasia of the scapula and pelvis (Lausch et al., 2008). A mouse model, droopy ear, has been described and consists of cranial malformations including misshapen and rotated ears, skeletal defects and reduced pigmentation (Candille et al., 2004; Singh et al., 2005; Farin et al., 2007; Lausch et al., 2008). In mice, *tbx15* is a transcriptional repressor and has a characterized binding site (Begemann et al., 2002; Farin et al., 2007). Little is known about the functions of *tbx15* in zebrafish, though sequence analysis and expression locations suggest that it is orthologous to human and murine *tbx15* (Begemann et al., 2002).

These two transcription factors are of interest due to microarray studies in which they are implicated in the hair cell regeneration process in chickens (Hawkins et al., 2007) and zebrafish (Liang, personal communication) as well as enriched in the zebrafish pLL (Gallardo et al., 2010). The functions of these genes and a known downstream target of Tbx5, *camK2b2*, on pLL development have been investigated. These studies identify a significant and previously unknown role for *tbx5* in the migration of the pLL primordium and subsequent development of the zebrafish pLL, at least in part via the downstream target *camK2b2*, and demonstrate that *tbx15* is important in modulating Tbx5 activity.

## Materials and Methods

### Zebrafish strains and handling

Animal care and husbandry were performed according to methods described by Westerfield et al. and in compliance with the National Human Genome Research Institute Animal Care and Use Committee guidelines (protocol # G-01-3) (Westerfield et al., 1986). All zebrafish embryos were incubated at 28°C. The Tg(−8.0cldnb:lynEGFP) transgenic strain was used for all experiments. Matings were performed in breeding tanks, and embryos for injection were collected approximately 20 minutes after males and females were mixed. In all cases, the point at which fish are combined in a breeding tank is taken to be time zero.

### Morpholino Injections

Two anti-sense morpholino oligonucleotides (morpholinos) were used to demonstrate phenotypes for each gene. For *tbx5*, we used the translation-blocking morpholino 5’ GGTGTCTTCACTGTCCGCCATGTCG and the splice-blocking morpholino ‘5 ACCGTGAGGCCTAAATACAGAAACA. For *tbx15*, we used the translation-blocking morpholino 5’ CAGCGGACCGTCTCCTGTCACTCAT and the splice-blocking morpholino 5’ TGGCCTCTGAATCTACAAAGAACAA. For *camK2b2*, we used the morpholinos previously described (Rothschild et al., 2009). A p53 blocking morpholino was also used as a control (Bill et al., 2009). Unless otherwise noted, data presented for Tbx5 morphants result from the splice-blocking morpholino; and data presented for Tbx15 morphants result from the translation-blocking morpholino. All morpholinos were injected into Tg(−8.0cldnb:lynEGFP) transgenic embryos at the 1-2 cell stage. Concentrations ranged from 250-1000 µM. Dual morphants received both *tbx5* and *tbx15* morpholinos at 750-1000 µM. Data were collected at room temperature using a Zeiss Axiovert 200M inverted fluorescent microscope. For each set of conditions, data were collected and pooled for analysis for wild type and morphant embryos from at least three different clutches.

### CaM-KII Inhibition

Calmodulin Kinase IINtide (Calbiochem #208920) was prepared according to the manufacturer’s instructions at a stock concentrations of 1 mM by dissolving in 100% DMSO. Dechorionated *cldnb:gfp* embryos were incubated from 20–21 hpf to 48 hpf in 50 µM CaM-KIINtide in E3 medium. Negative control embryos were treated with E3 containing 1% DMSO.

### Rescue Construct

The CamKII rescue construct used for rescue of Tbx5 and Tbx15 morphant phenotypes is a linearized plasmid expression construct containing the human ∂C CamKII variant previously described and kindly provided by the Tombes laboratory (Rothschild et al., 2009). The rescue construct was co-injected with morpholino or alone at a concentration of 40 ng/uL.

### In situ hybridization

*In situ* hybridization was performed on wild type embryos and *tbx5, tbx15* and *camk2b2* morphant embryos that were fixed in 4% paraformaldehyde at 36 hpf. *In situ* hybridization was carried out according to standard protocols. Staining times varied according to the efficiency of the digoxigenin-labeled riboprobes, but generally required overnight staining at room temperature. Riboprobes for mRNA from *tbx5, tbx15, cxcr7b, cxcr4b, fgf10a* and *notch3* were made using linearized plasmid cDNAs obtained from the Open Biosystems clones: 6967993, 7065736, 8746628, 7238701, 7230079, and 7995007, respectively. Linearized clones were used to make cDNA by PCR with T7 extended antisense primers, which was then *in vitro* transcribed with the Roche DIG RNA Labeling Kit (SP6/T7). Imaging was done using a Zeiss Stemi SV 11 dissecting scope and OpenLab 3.1.5 software. *camk2b2* riboprobes were made using two primer sets against the *camk2b2* cDNA sequence, an outside set and a nested set, as a cDNA clone is not presently available for this gene.

### Immunofluorescent Staining

Antibody stains were performed on Tg(−8.0cldnb:lynEGFP) control and morphant embryos using anti-GFP-FITC conjugated antibody. Confocal images were acquired at room temperature using a Zeiss LSM 510 NLO Meta system mounted on a Zeiss Axiovert 200M microscope with a 5x or a 10x objective using the Zeiss AIM 4.0 v2 software package. Excitation wavelengths of 488nm were used for GFP or FITC detection. Fluorescent emissions were collected in band-pass filters 500-550nm IR blocked, 575-615nm, 641-705nm collected in the Meta detector and 390-465nm IR blocked, respectively. All pinholes were set at 1.00 Airy units (AU), which corresponds to an optical slice of 0.8µm. All confocal images were of frame size 512 pixels by 512 pixels, scan zoom of 1 and line averaged 4 times.

### Quantification of expression levels

qRT-PCR experiments were performed from total RNA isolated from control and morphant embryos using Invitrogen’s 2-step sybr-greenER qRT-PCR kit for iCycler (Bio-Rad). Primers for each gene tested were designed using PrimerBLAST (www.ncbi.nlm.nih.gov/tools/primer-blast) to ensure specificity. All results were normalized using ß-actin primers.

### Data Analysis and Statistics

Statistical analysis of neuromast count data was done using Microsoft Excel t-test functions. All t-tests were 2-tailed, unpaired, and assuming unequal variance. Analysis of phenotype data was done using Fisher’s exact test. In all cases, sample sizes (n) were reported per fish and not per pLL. All statistical analyses were done based on the total number of neuromasts per fish. qRT-PCR analysis was done using the standard 2^∆∆cT^ equation. Each sample was run in at least triplicate.

## Results

### Tbx5 morphant phenotype

To investigate the role of *tbx5* in zebrafish pLL development, morpholinos against Tbx5 RNA were injected into Tg(−8.0cldnb:lynEGFP) embryos. Morphant embryos were raised to 48 hpf and examined for pLL effects. Heart chambers in the morphants failed to loop, resulting in cardioedemas, suggesting that the heartstrings mutant had successfully been phenocopied in the morphants (Fig. 1A). Furthermore, morphants did not produce full pectoral fins, though fin bud development was initiated (data not shown), which is consistent with the previously described morphant phenotype (Garrity et al., 2002). Additional analysis showed that two different morpholinos against *tbx5* but not a negative control morpholino resulted in reduction of the average number of neuromasts in the morphant pLL compared with the wild type (Fig. 1B). Quantification demonstrates that the average number of neuromasts per pLL is reduced from 8 in the wild type to 2 in the morphants, which is a statistically significant reduction (Fig. 1C).

**Figure 1.**
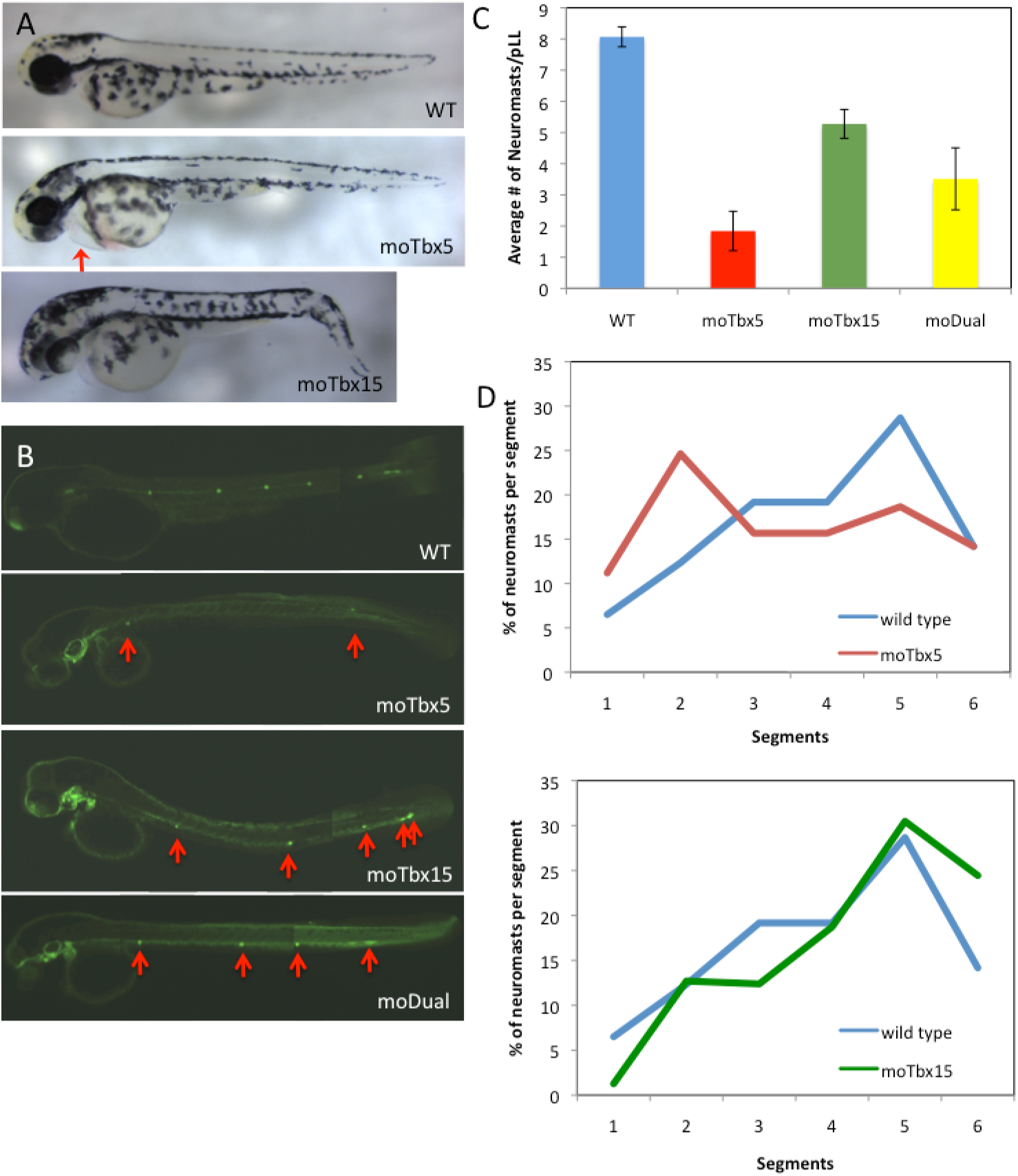
Tbx5 and Tbx15 morphants had reduced numbers of neuromasts and altered neuromast spacing while dual morphants demonstrated an intermediate phenotype. (A) The Tbx5 morphant phenotype is consistent with the published *tbx5* mutant phenotype, *heartstrings*, and demonstrates reduced pectoral fin buds with failed pectoral fin development (not pictured), edema in the heart chambers (arrow) following failed looping to form the atrium and ventricle, and failure to survive to adulthood. The Tbx15 morphant phenotype is relatively mild, although morphant embryos frequently have a bent tail. (B) Neuromast deposition in the pLL was analyzed in wild type, Tbx5, Tbx15 and Dual (Tbx5 + Tbx15) morphant Tg(−8.0cldnb:lynEGFP) embryos at 48 hpf. Two morpholinos against *tbx5* were utilized, a transcription-blocking morpholino (data not shown) and a splice-blocking morpholino. Two morpholinos against *tbx15* were utilized, a transcription-blocking morpholino and a splice-blocking morpholino (data not shown). All morpholinos against *tbx5* and *tbx15*, but not a standard control, produced fewer numbers of pLL neuromasts (arrows). Tbx5 and Tbx15 morpholinos were co-injected and neuromast deposition was analyzed in the resulting dual morphant embryos at 48 hpf. The dual morphants demonstrated a more severe phenotype than the Tbx15 morphants but a less severe phenotype than the Tbx5 morphants. (C) Neuromasts quantification shows a significant reduction in number of pLL neuromasts in both Tbx5 and Tbx15 morphants compared with the wild type (n(WT) = 53; n(moTbx5) = 38, p < 0.05; n(moTbx15) = 59, p < 0.05). The p53 control did not produce a significant reduction in average number of pLL neuromasts (not shown). Quantification of the dual morphant neuromasts demonstrates a significant reduction in average neuromast count compared with the Tbx15 morphant (n(moTbx15) = 59, n(moDual) = 36, p < 0.05). A significant increase in average neuromast count is noted in the dual morphant over the Tbx5 morphant (n(moTbx5) = 38, n(moDual) = 36, p < 0.05) meaning the double phenotype is intermediate to the two morphants alone. Data are reported as means +/− 95% confidence intervals. (D) Neuromast positioning was analyzed by dividing the fish body into six equal segments from the posterior edge of the otic placode to the tip of the tail and counting the neuromasts per segment. In Tbx5 morphants, neuromasts are located with the highest frequency in segment 2, whereas neuromast frequency is highest in segment 5 in the wildtype (n(WT) = 36, n(moTbx5) = 25). The frequency of deposition differs for the Tbx5 morphant significantly in segments 1, 2 and 5 (p < 0.05). Tbx15 morphants demonstrate an overall posterior shift in neuromast deposition. moTbx15 neuromast frequency differs significantly from wild type frequency in segments 1, 3, and 6 (n(WT) = 36, n(moTbx15) = 30, p < 0.05).

Furthermore, a pattern in neuromast spacing was observed in the morphant pLL. To document this pattern, embryos were divided into six equal segments from the back edge of the otic vesicle to the tip of the tail, and the neuromasts located in each segment were counted. The largest neuromast concentration in the morphants was found in segment 2 whereas neuromasts were spread more evenly in wild type embryos at 48 hpf with a modest peak in the 5^th^ segment (Fig. 1D). Thus, an early deposition pattern in addition to an overall reduction in neuromast number was present in Tbx5 morphants.

### Tbx15 morphant phenotype

Given that *tbx15* is upregulated in the same regenerative and developmental processes as tbx5, the role of *tbx15* in pLL development was also investigated. Morpholinos against Tbx15 mRNA were injected into Tg(−8.0cldnb:lynEGFP) embryos. Embryos were raised to 48 hpf and examined. Morphant embryos appeared generally normal, though some possessed a bend in the tail (Fig. 1A). Two morpholinos against *tbx15*, but not a control morpholino, produced a reduction in the number of neuromasts deposited in the pLL (Fig. 1B). Quantification reveals a statistically significant reduction in the average number of neuromasts from 8 in wild type to about 5 in morphants (Fig. 1C).

As with Tbx5 morphants, a pattern in neuromast spacing was noted and quantified in Tbx15 morphants. In contrast to the Tbx5 pattern, however, Tbx15 morphants possessed a posterior shift in neuromast deposition, as evidenced by a significant reduction in neuromast frequency in segments 2 and 3 while neuromast frequency is significantly increased in segment 6, when compared with wild type (Fig. 1D). In Tbx15 morphants, a decrease in neuromast count is demonstrated (though this decrease is not as severe as that for Tbx5 morphants) along with a posterior shift in neuromast deposition.

Morpholinos have been shown to activate the p53 pathway in zebrafish, which can be alleviated by co-injection with morpholinos that target p53 (Bill et al., 2009). Neither morphant phenotype was significantly affected by co-injection with p53 morpholinos (data not shown).

### tbx5 and tbx15 are expressed in the developing pLL

Having found that both *tbx5* and *tbx15* produce a significant pLL phenotype we determined the expression locations for each gene. mRNA-specific probes for *tbx5* and *tbx15* were used for *in situ* hybridization in Tg(−8.0cldnb:lynEGFP) embryos. Embryos were fixed at 36 hpf to allow visualization of expression in the migrating primordium. Enhanced staining in the neuromasts, interneuromast cells, and the migrating primordium indicate that *tbx5* and *tbx15* are both expressed in the zebrafish pLL (Fig. S1).

### Dual morphants have an intermediate phenotype

Given that T-box transcription factors can function as monomers or as dimers, it was essential to establish evidence of interaction between Tbx5 and Tbx15 in the pLL developmental process. Thus, morpholinos for *tbx5* and *tbx15* were co-injected to create dual morphants. Dual morphant zebrafish embryos averaged 3 – 4 neuromasts per pLL at 48 hpf, which is an intermediate level between that of Tbx5 morphant and Tbx15 morphant phenotypes (Fig. 1B,C), indicating that knocking down Tbx15 partially rescues the Tbx5 morpholino phenotype. The intermediate phenotype along with the differences in neuromast positioning between Tbx5 and Tbx15 morphants suggest that the two transcription factors act in the same pathway to influence pLL development but that they function in an opposing manner.

### camK2b2 knock-down produces pLL defects similar to the Tbx5 morphant phenotype

The specific roles of *tbx5* and *tbx15* in pLL development were investigated by examining the effects of a known downstream target of Tbx5 in zebrafish. Calcium/calmodulin protein kinase II, ß2 (*camK2b2*) acts downstream of Tbx5 in development of both the pectoral fins and the heart and may be a direct transcriptional target of Tbx5 (Rothschild et al., 2009). CamK2b2 activity in zebrafish pLL development was elucidated by injecting single-cell stage embryos with *camK2b2* morpholinos. Morphant embryos were examined at 48 hpf. Two *camK2b2* morpholinos but not a control morpholino resulted in pLL defects. Morphants demonstrate a significant reduction in the average number of neuromasts per pLL compared to wild type (Fig. 2A,B). This reduction is similar to but not as severe as the reduction attributable to *tbx5* knockdown.

**Figure 2.**
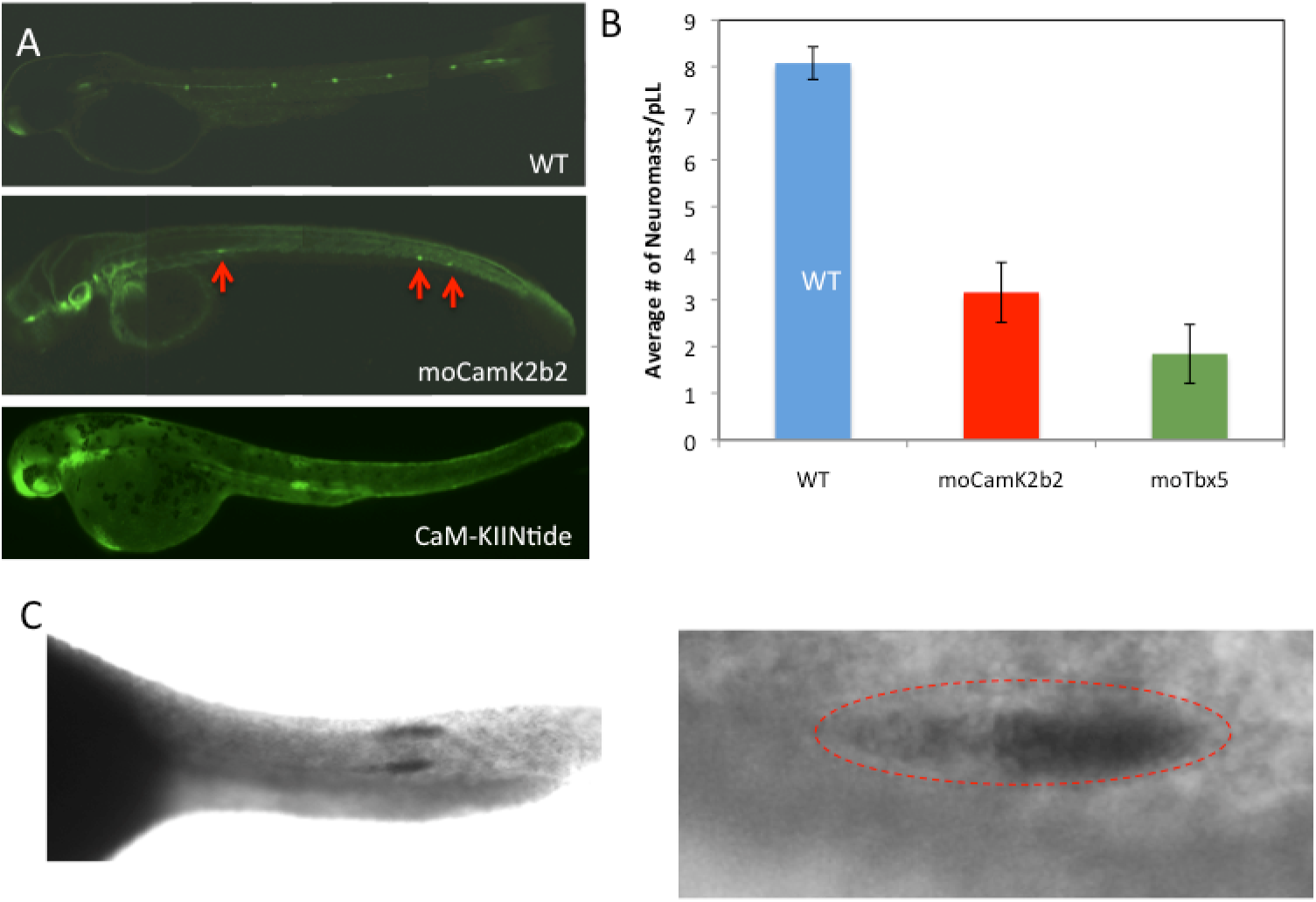
CamK2b2 knockdown and CaM-KII inhibition result in pLL defects. *camk2b2* is expressed in the developing pLL. (A) Neuromast deposition in the pLL was analyzed in wild type, Tbx5 morphant, and CamK2b2 morphant cldnb:gfp embryos at 48 hpf. Two different morpholinos against *camk2b2* were utilized. Both morpholinos, but not a p53 co-injected control, produced reduced numbers of pLL neuromasts, consistent with the Tbx5 morphant phenotype. Defects in pLL primordium migration and neuromast deposition also resulted following treatment with the CaM-KII inhibitor CaM-KIINtide. (B) Neuromasts quantification shows a significant reduction in the number of pLL neuromasts compared with the wildtype (n(WT) = 53, n(moCamK2b2) = 34, p < 0.05), though the reduction is not as severe as that seen in the Tbx5 morphants. Data are reported as means +/− 95% confidence intervals. (C) *In situ* hybridization with digoxigenin-labeled RNA probes against CamK2b2 mRNA demonstrates expression of *camk2b2* in the developing pLL.

Corroborating evidence of camK2b2 impacts on pLL development were obtained using the CaM-KII inhibitor CaM-KIINtide (Chang et al., 1998). Inhibition of activated CamKII resulted in arrested primordium migration and reduced neuromast deposition (Fig. 2A). To verify that *camk2b2* is expressed in the pLL, riboprobes against Camk2b2 mRNA were used for *in situ* hybridization in Tg(−8.0cldnb:lynEGFP) embryos fixed at 36 hpf. Hybridization results demonstrated that *camk2b2* is expressed in the pLL and the migrating pLL primordium (Fig. 2C).

### Ectopic expression of CamKII partially rescues the Tbx5 morphant phenotype and exacerbates the Tbx15 morphant phenotype

Having shown that knockdown of *camk2b2* leads to reduced neuromast count and that *camk2b2* is expressed in the pLL, the impact of ectopic expression of CamKII on Tbx5 and Tbx15 morphant phenotypes was examined. A linearized plasmid construct containing the human ∂C variant of CamKII driven by a constitutive promoter was co-injected with either *tbx5* or *tbx15* morpholinos as previously described (Rothschild et al., 2009). Co-injection with the Tbx5 morpholino results in an increase in the number of neuromasts in each pLL compared with the Tbx5 morpholino alone (Fig. 3A). This increase is a statistically significant (p < 0.05) improvement in the Tbx5 morphant phenotype, indicating that replacing some CamKII function partially rescues the Tbx5 morphant phenotype in the zebrafish pLL (Fig. 3B), even if this expression is ectopic rather than spatially restricted. Furthermore, co-injection of the rescue construct with the *tbx15* morpholino results in significantly fewer neuromasts per pLL (Fig. 3C,D) compared to *tbx15* morpholinos alone. Injecting the CamKII expression construct alone does not significantly impact pLL development, so ectopic expression alone is not harmful, which is not surprising given that CamKII must be activated by calcium signaling in order to be effective. That being the case, the exacerbated phenotype noted when CamKII is ectopically expressed in Tbx15 morphants is specifically attributable to increased CamKII activity in a background of knocked-down *tbx15* activity.

**Figure 3.**
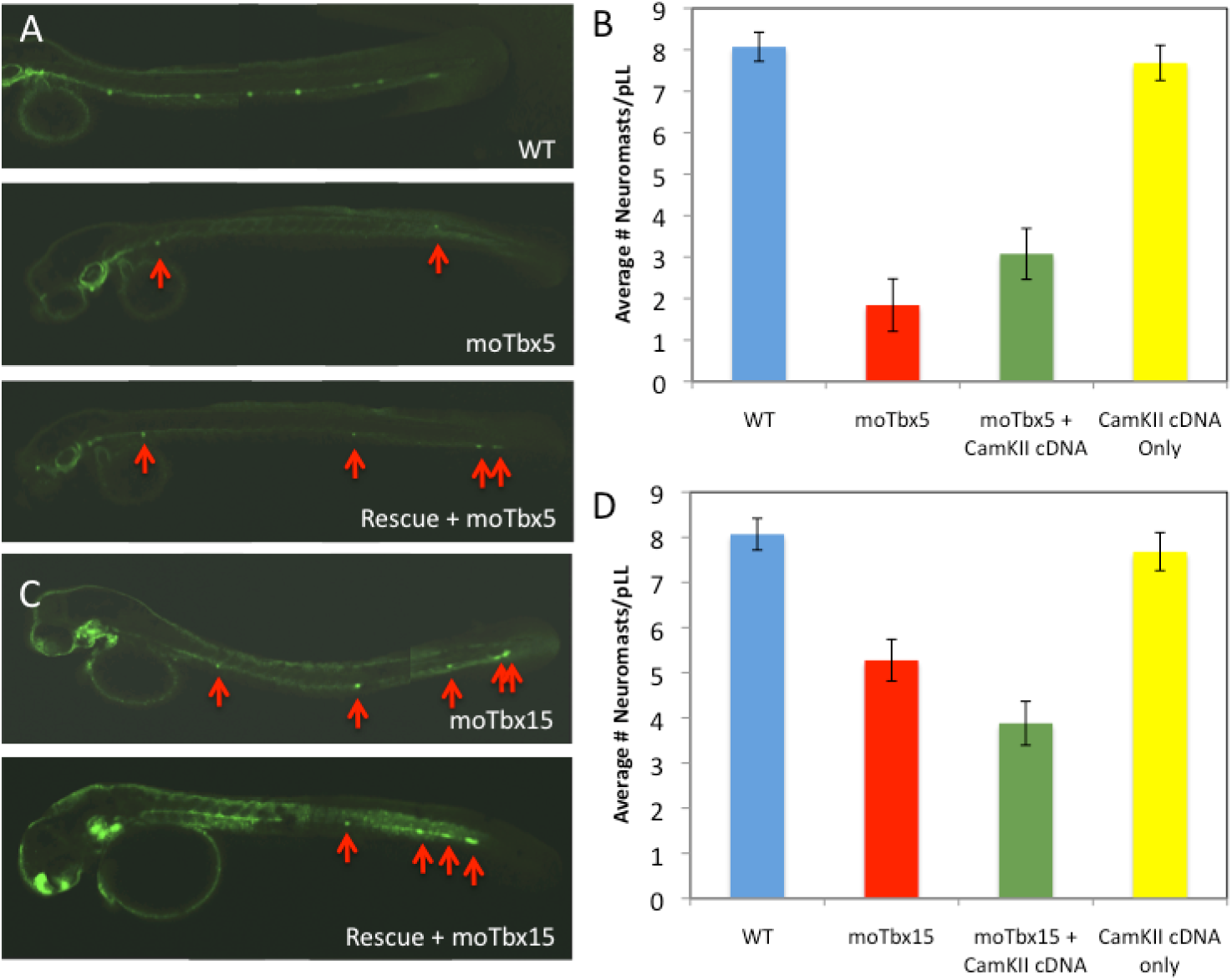
Ectopic CamKII partially rescues the Tbx5 morphant phenotype and exacerbates the Tbx15 phenotype (A) Tbx5 morpholinos were co-injected with a linearized DNA expression construct for the human δc CamKII variant. Neuromast deposition was analyzed in the morphants, revealing an improved pLL phenotype. (B) Quantification of the neuromasts demonstrated a significant increase in the average number of neuromasts per pLL in the rescued Tbx5 morphants in comparison with the unrescued Tbx5 morphants (n(moTbx5) = 38, n(moTbx5 + CamKII cDNA) = 50, p < 0.05). The average number of neuromasts per pLL, however, is not restored to or near to the wild type, thus the rescue is partial. Data are reported as means +/− 95% confidence intervals. (C) Tbx15 morpholinos were co-injected with a linearized DNA expression construct for the human δc CamKII variant. Neuromast deposition was analyzed in the morphants, revealing a more severe pLL phenotype than Tbx15 morphants alone. Data are reported as means +/− 95% confidence intervals. (D) Neuromasts quantification demonstrated a significant decrease in the average number of neuromasts per pLL in the rescued Tbx15 morphants in comparison with the unrescued Tbx15 morphants (n(moTbx15) = 59, n(moTbx5 + CamKII cDNA) = 79, p < 0.05). Injection of rescue construct alone resulted in no significant change in average number of neuromasts. Data are reported as means +/− 95% confidence intervals.

### camK2b2 expression is lost in Tbx5 morphants but not in Tbx15 morphants

The rescue results support the hypothesis that *camk2b2* is a downstream target of Tbx5 in the developing pLL, as in heart and pectoral fin development. Therefore, *camk2b2* expression was analyzed in Tbx5 morphant embryos at 36 hpf by in situ hybridization and is lost in the pLL and primordium of these fish (Fig. S2). Additionally, qRT-PCR was used to quantify the effect of *tbx5* knockdown on *camk2b2* expression. In multiple replicates, *camk2b2* mRNA was not detectable in total RNA extracts from Tbx5 morphants (Fig. 4A), though it was detectable in wild type and all other morphant conditions (Fig. 4B,C,D). Taken together, these data demonstrate a loss of *camk2b2* expression in Tbx5 morphants and suggest that *tbx5* is the primary *camk2b2* activator in all early zebrafish embryo tissues.

**Figure 4.**
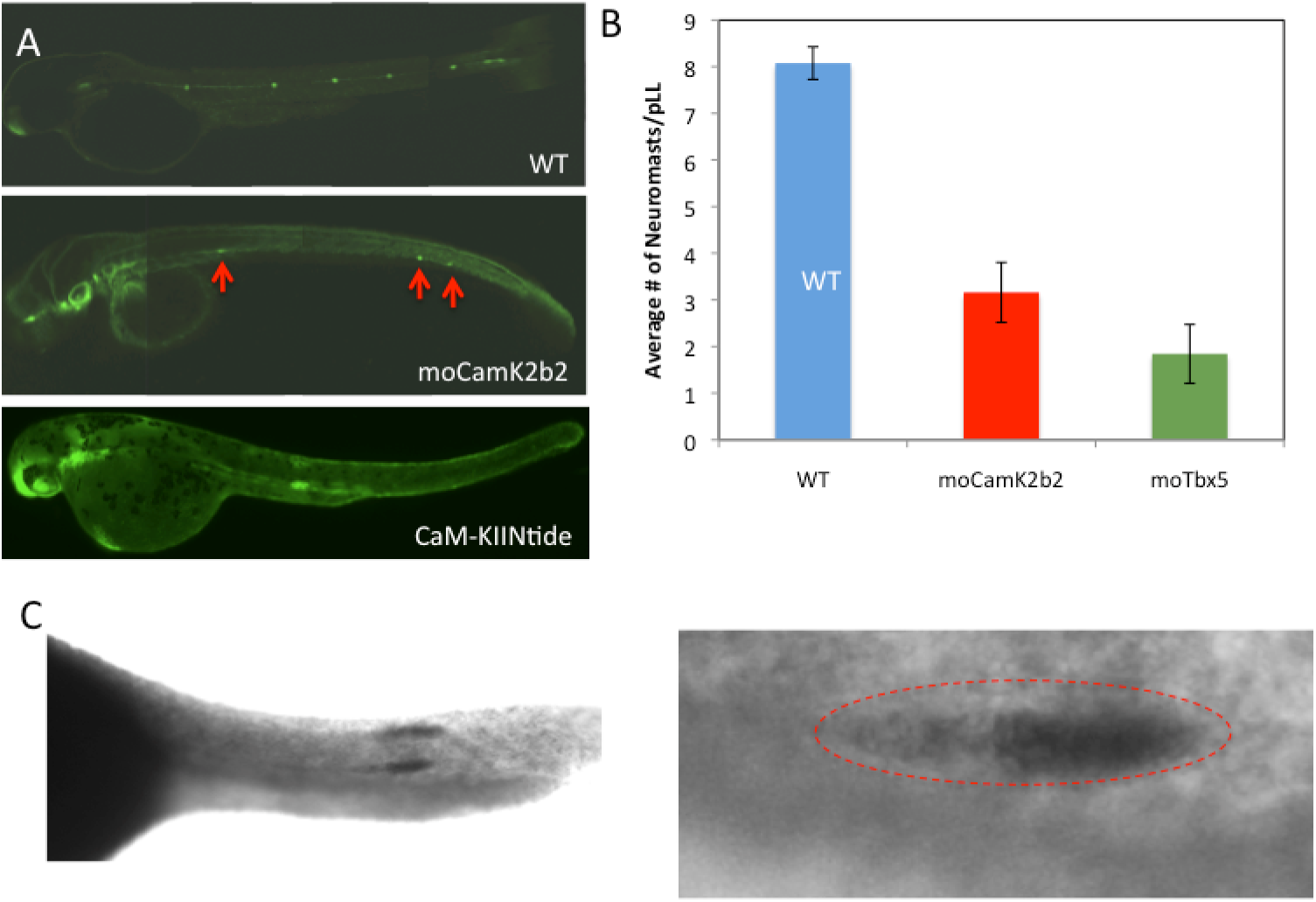
Expression levels of *tbx5, tbx15* and *camK2b2* are altered in morphant embryos. (A) qRT-PCR analysis shows that, following treatment with the splice-blocking morpholino against *tbx5*, Tbx5 and Tbx15 mRNA levels are both significantly reduced (p(tbx5) < 0.05, p(tbx15) < 0.05) while CamK2b2 mRNA is not detected. (B) qRT-PCR analysis following treatment with the translation-blocking morpholino against *tbx15* indicate that loss of Tbx15 results in significant upregulation of *tbx15* (p < 0.05) and *tbx5* (p < 0.05). *camK2b2* expression was not significantly impacted. (C) Camk2b2 morphants were analyzed by qRT-PCR. Data indicate that *tbx5, tbx15*, and *camk2b2* are all significantly upregulated following loss of Camk2b2 (p(tbx5) < 0.05, p(tbx15) < 0.05, p(camk2b2) < 0.05). (D) Embryos injected with the rescue construct were analyzed by qRT-PCR. Neither *tbx5* nor *camk2b2* expression was significantly impacted; however, *tbx15* was upregulated following ectopic expression of CamKII (p < 0.05). Data represent means +/− SEM.

Similarly, the impact of *tbx15* knockdown on *camk2b2* was examined by *in situ* hybridization and qRT-PCR. In this case, it was hypothesized that there would be no significant impact on *camk2b2* expression, as previous data do not suggest that *camk2b2* is a downstream target of Tbx15. As expected, *camk2b2* expression is detected by staining of the pLL and primordium (Fig. S2). Furthermore, qRT-PCR analysis demonstrates that Camk2b2 mRNA levels are not significantly impacted by *tbx15* knockdown (Fig. 4B).

### Expression levels of tbx5, tbx15 and camK2b2 are altered in morphant embryos

The data suggest that *tbx5* and *tbx15* are involved in the same pathway affecting pLL development and that they function in opposing directions within this pathway. Furthermore, it appears that *camk2b2* is a downstream target of Tbx5, but not of Tbx15 in this process. To investigate the interactions that occur amongst these genes, expression of each was analyzed in all three morphants by qRT-PCR. In Tbx5 morphants, *tbx5* and *tbx15* are both significantly downregulated while *camk2b2* is completely eliminated (Fig. 4A). Downregulation of *tbx5* is expected, as a splice-blocking morpholino that would result in nonsense-mediated decay was used to make these morphants. In Tbx15 morphants, *tbx15* and *tbx5* were both upregulated (Fig. 4B). In Camk2b2 morphants, expression of all three genes is significantly upregulated (Fig. 4C). To determine if *tbx15* expression is responsive to CamKII levels, the CamKII expression construct used for rescue was injected alone into Tg(−8.0cldnb:lynEGFP) embryos. qRT-PCR analysis of total RNA from these embryos at 36hpf demonstrated that *tbx5* and *camk2b2* expression were not affected, but *tbx15* was upregulated (Fig. 4D). Based on these data, a model is proposed in which *tbx5* activates *camk2b2* expression, and in parallel activates *tbx15*. Tbx15 is then important in negatively modulating *tbx5* in combination with *camk2b2*, as evidenced by upregulation of *tbx5* in Tbx15 morphants and in Camk2b2 morphants. This feedback loop, then, is essential for development of a normal pLL (Fig. 7).

### Primordium patterning is disrupted in morphant embryos

Visible defects in primordium shape were observed in morphant embryos, thus we hypothesized that *tbx5* and *tbx15* are involved in primordium patterning. In normal Tg(−8.0cldnb:lynEGFP) embryos, primordia are elongated cell masses with bright focal points corresponding to the centers of rosette structures. In Tbx5 and Camk2b2 morphant primordia, these focal points are frequently lost, indicating that rosettes are not properly forming (Fig. S3). In Tbx15 morphants, there was a reduction in rosettes numbers, though defects were not as extreme as those seen in Tbx5 and Camk2b2 morphants. Furthermore, primordia deformations were documented in all morphants, including fragmentation and loss of overall primordium structure (Fig. S3). Other defects noted in Tbx5 morphants include occasional failed migration and migration along alternate paths similar to mutations in *cxcr4b* (David et al., 2002).

### Expression of primordium patterning markers is affected in morphants

Based on observations of primordium defects, the specific impacts of *tbx5* and *tbx15* on markers of primordium patterning were investigated. The first markers chosen were *cxcr4b* and *cxcr7b*, as their roles in maintenance of primordium integrity and in migration have been extensively studied. In addition, many of the primordium defects noted are similar to those seen in Cxcr7b morphants (Valentin et al., 2007). To examine these markers, *in situ* hybridization was performed in Tbx5, Tbx15 and Camk2b2 morphants using *cxcr4b* and *cxcr7b* riboprobes. Expression of *cxcr4b* is expected mainly in the leading half of the primordium, as in the wild type (93%, n = 29). This expression pattern is maintained in both Tbx15 (69%, n = 26) and Camk2b2 (100%, n = 12) morphants. Interestingly, expression of *cxcr4b* is expanded to the entire primordium in Tbx5 morphants (Fig. 5A) (70%, n = 30). Expression of *cxcr7b* is expected in the trailing half of the primordium. This pattern is seen in wild type (94%, n = 34) and Tbx15 (69%, n = 34), Camk2b2 (82%, n = 29), and Tbx5 morphants (74%, n = 17) (Fig. 5A).

**Figure 5.**
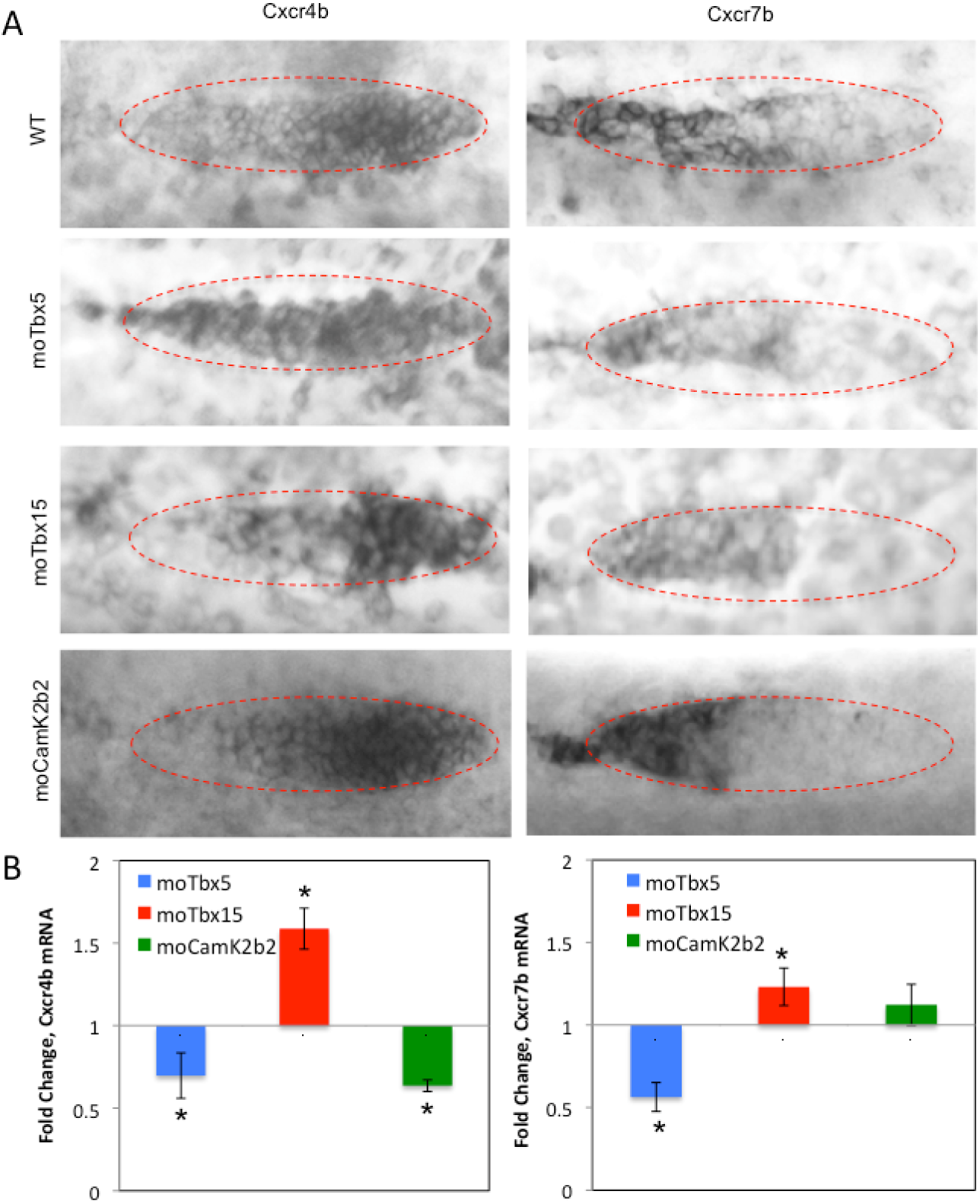
*cxcr4b* and *cxcr7b* have altered expression patterns and levels in Tbx5 morphants. (A) In situ hybridization was performed on WT, Tbx5, Tbx15, CamK2b2 morphant Tg(−8.0cldnb:lynEGFP) embryos at 36 hpf using riboprobes against *cxcr4b* and *cxcr7b* mRNA. Primordium location is indicated by the dotted oval in all images. In Tbx5 morphants, *cxcr4b* expression is expanded from the leading zone of the primordium in the wild-type case encompassing the trailing zone as well in the Tbx5 morphants. The expression patterns of *cxcr4b* and *cxcr7b* are unchanged in Tbx15 and CamK2b2 morphants. (B) qRT-PCR analysis of WT, Tbx5, Tbx15, and CamK2b2 morphant embryos at 36 hpf for Cxcr4b and Cxcr7b mRNA. *cxcr4b* expression is significantly reduced in both Tbx5 and Camk2b2 morphants (p(tbx5) < 0.05, p(camk2b2) < 0.05) while it is upregulated in Tbx15 morphants (p(tbx15) < 0.05). *cxcr7b* expression is likewise reduced in Tbx5 morphants (p(tbx5) < 0.05) and is upregulated in Tbx15 morphants (p(tbx15) < 0.05). Expression of *cxcr7b* is not significantly altered in CamK2b2 morphants. Data represent means +/− SEM.

Additionally, qRT-PCR was used to examine expression levels of both *cxcr4b* and *cxcr7b* in wild type and morphants. In both Tbx5 and Camk2b2 morphants, *cxcr4b* is downregulated while it is upregulated in Tbx15 morphants (Fig. 5B). These data suggest that both *tbx5* and *camk2b2* act upstream of *cxcr4b* and that the *tbx5/tbx15* negative feedback loop is involved in determining *cxcr4b* expression levels. Furthermore, it was determined that, like *cxcr4b, cxcr7b* is downregulated in Tbx5 morphants (Fig. 5B). However, knockdown of *camk2b2* does not impact *cxcr7b* expression, indicating that Camk2b2 is involved in the expression of *cxcr4b* but not *cxcr7b*. Finally, it was noted that knockdown of *tbx15* results in upregulation of *cxcr7b* (Fig. 5B), indicating that this negative feedback loop may be important in establishing proper *cxcr4b* and *cxcr7b* expression.

*In situ* hybridization was performed for two additional primordium markers: *fgf10a*, a potential indicator of Fgf signaling activity and *notch3*, a marker of rosette patterning. Normally, *fgf10a* is expressed in the leading half of the primordium and in rosette centers in the trailing half of the primordium. This pattern of *fgf10a* expression was observed in the wild type embryos (96%, n = 29) and in Tbx15 (74%, n = 26) and Camk2b2 (100%, n = 12) morphant embryos but not in Tbx5 morphant embryos (Fig 6A). Here, *fgf10a* expression is shifted to the trailing edge of the primordium (67%, n = 30), indicating that *tbx5* knockdown interferes with *fgf10a* expression and possibly FGF signaling. In the case of *notch3*, normal expression yields staining throughout the primordium with the exception of rosette centers, as *notch3* has previously been shown to play a role in limiting expression of *deltaA* and *atoh1a* to the rosette centers, which is important in committing precursor cells to a hair cell fate (Millimaki et al., 2007). The normal staining pattern was exhibited in wild type (100%, n = 34), Tbx15 morphants (73%, n = 34) and Camk2b2 morphants (92%, n = 29); however, expression was altered in Tbx5 morphants such that *notch3* staining was clearly visible in rosette centers (Fig. 6A) (66%, n = 17). Thus, the data further suggest that *tbx5* is involved in rosette patterning. qRT-PCR was used to examine expression levels of *fgf10a* and *notch3* in Tbx5, Tbx15 and CamK2b2 morphants. Two additional markers, *atoh1a* and *lef1*, were studied. *atoh1a* expression specifies cells fated to become hair cells and is involved in the hair cell development and maturation process. Lef1 is a transcription factor that is activated by Wnt/ß-catenin signaling and is implicated in numerous developmental processes. All four markers of primordium patterning are downregulated in Tbx5 morphants. *notch3* and *atoh1a* are also downregulated in Camk2b2 morphants (Figs. 6B, S4), suggesting that *tbx5* mediates expression of *notch3* via *camk2b2* and that its impact on expression of *fgf10a* is independent of *camk2b2*. In all cases, knockdown of *tbx15* does not significantly impact marker expression (Figs. 6B, S4).

**Figure 6.**
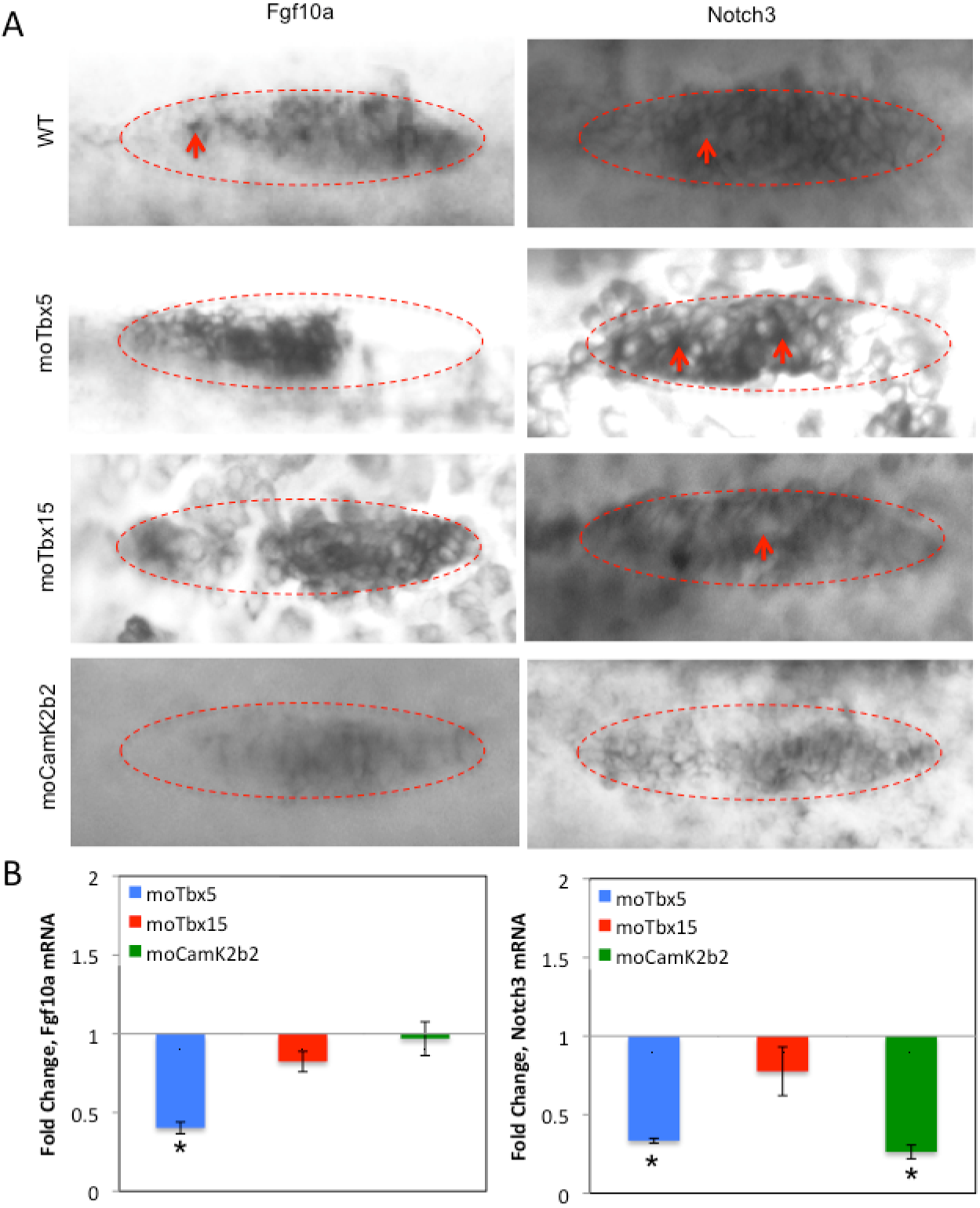
Fgf10a and Notch3 have altered expression patterns in Tbx5 morphants. (A) In situ hybridization was performed on WT, Tbx5, Tbx15, and CamK2b2 morphant cldnb:gfp embryos at 36 hpf using riboprobes against *fgf10a* and *notch3* mRNA. The primordium is indicated by the dotted oval in each image. In the wild type, *fgf10a* is expressed in the leading zone of the primordium and at rosette centers in the trailing zone (arrow), while *notch3* is expressed throughout the primordium with the exception of rosette centers (arrow). The wild type pattern for both *fgf10a* and *notch3* is observed in Tbx15 and Camk2b2 morphants. However, *fgf10a* expression in Tbx5 morphants is shifted from the leading zone of the primordium to the trailing zone. In addition, *notch3* expression in Tbx5 morphants is noted at rosette centers (arrows), indicating that both *fgf10a* and *notch3* expression patterns are altered in Tbx5 morphants. (B) qRT-PCR analysis was conducted on WT, Tbx5, Tbx15 and Camk2b2 morphants at 36 hpf for Fgf10a and Notch3 mRNA. *fgf10a* expression is significantly reduced in Tbx5 morphants (p(tbx5) < 0.05) but is not significantly altered in Tbx15 and Camk2b2 morphants. *notch3* expression is significantly reduced in both Tbx5 (p(tbx5) < 0.05) and Camk2b2 (p(camk2b2) < 0.05) morphants. *notch3* expression is not significantly changed in Tbx15 morphants. Data represent means +/− SEM.

## Discussion

Our knowledge of the requirements for successful zebrafish pLL development is dependent upon an ever-increasing list of genes. Disrupting any of these genes results in defects in primordium tissue patterning, cell proliferation, neuromast deposition, and/or migration. Furthermore, these processes are coupled such that a deficiency in patterning, for example, may lead to deposition or migratory defects (Lecaudey et al., 2008). Signaling molecules active within the primordium are vitally important in this developmental process. In particular, the Wnt/FGF balance is essential for establishing localization and appropriate gene expression within the primordium, allowing undifferendiated cells at the leading edge to undergo morphological changes, apico-basal polarization, and apical constriction, leading to formation of radial rosette structures (Lecaudey et al., 2008; Nechiporuk and Raible, 2008; Aman and Piotrowski, 2009; Ma and Raible, 2009). Within these rosettes, central foci are marked by *fgf10a, deltaA* and *atoh1a* expression while *notch3* expression toward the rosette periphery appears to be key in restricting expression of *deltaA*, and subsequently *atoh1a*, to cells that eventually become functioning hair cells of the mature neuromast (Itoh and Chitnis, 2001; Millimaki et al., 2007; Nechiporuk and Raible, 2008). Furthermore, differential expression of *cxcr4b* and *cxcr7b* in the primordium is required for migration, an important process in ensuring that neuromasts are distributed along the body of the fish (Haas and Gilmour, 2006; Dambly-Chaudiere et al., 2007; Valentin et al., 2007; Aman and Piotrowski, 2008). Yet major gaps in knowledge prevent a broader understanding of the processes that govern pLL development in spite of numerous potential models.

To date, these models have not included T-box transcription factors. We demonstrate here that *tbx5* and *tbx15* are involved in zebrafish pLL development and that they participate in a negative feedback loop such that *tbx5* activity is maintained at a precise level, allowing for appropriate activation of downstream targets. These targets subsequently affect expression and localization of *cxcr4b, cxcr7b, fgf10a, notch3, lef1*, and potentially other genes important for pLL development. We further show that *camk2b2* is a downstream target of tbx5, is at least one upstream activator of tbx15 in the negative feedback loop, and plays a role in expression of *cxcr4b, notch3*, and *atoh1a* (Fig. 7).

**Figure 7.**
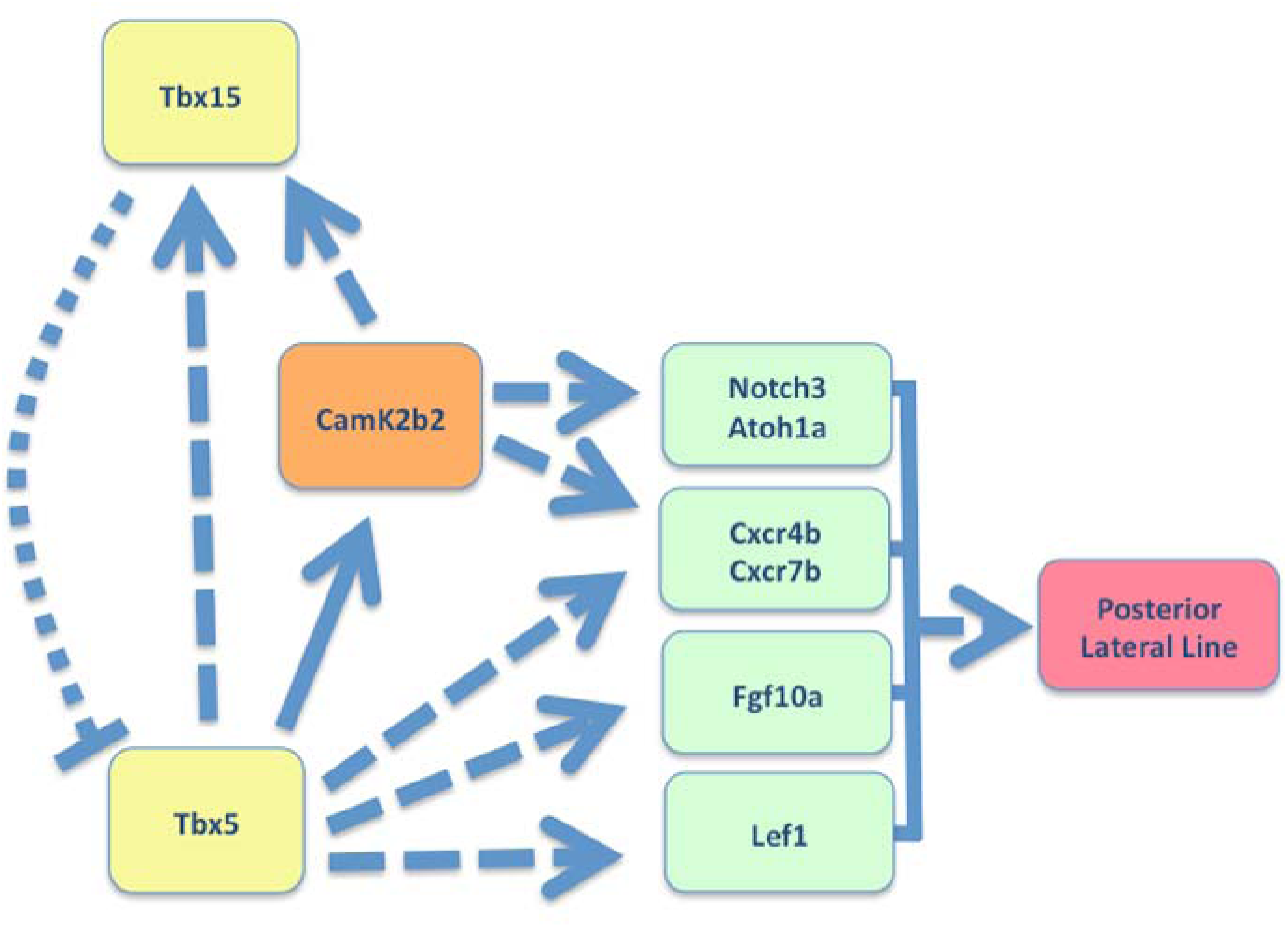
Proposed pathway for effects of *tbx5, tbx15* and *camk2b2* on pLL development. Tbx5 is a transcriptional activator of *camk2b2*. CamKII upregulates *tbx15*, which is a *tbx5* repressor. Thus these three participate in a negative feedback loop that modulates expression of both *tbx5* and *camk2b2*. Tbx5 also upregulates *notch3, atoh1a, cxcr4b, cxcr7b, fgf10a* and *lef1*. Some of these effects may be mediated through Camk2b2, as Camk2b2 also upregulates *notch3, atoh1a* and *cxcr4b*. *cxc4rb* is potentially affected by Tbx5 via two different mechanisms as Camk2b2 affects the expression level of *cxcr4b* but not the expression pattern, while Tbx5 morphants have an altered expression pattern in addition to an altered expression level. The genes affected implicate Tbx5 as an upstream activator of multiple signaling pathways that are involved in zebrafish pLL development.

Morphant analysis reveals disruption of pLL development following knockdown of either *tbx5* or *tbx15* and opposing shifts in deposition: an anterior shift in Tbx5 morphants and a posterior shift in Tbx15 morphants. This, along with partial rescue of the Tbx5 phenotype by knockdown of *tbx15* evidenced by the dual morphant intermediate phenotype, implicates these transcription factors in the same developmental pathway and suggests that they exert their effects in opposing directions. Camk2b2 morphants exhibit a phenotype similar to that of Tbx5 morphants. Data indicate that *camk2b2* transcripts are absent in Tbx5 morphants and that ectopic restoration of CamKII function is sufficient to provide partial rescue of the Tbx5 phenotype. Interestingly, ectopic “restoration” of CamKII function worsens the Tbx15 morphant phenotype. These results suggest that *camk2b2* activity is upregulated by Tbx5 and inhibited by Tbx15. However, further analysis by qRT-PCR demonstrates that Tbx15 is inhibiting activity at the level of *tbx5*, subsequently resulting in balanced expression of both *tbx5* and *camK2b2* (Fig. 7).

Analysis by qRT-PCR provides evidence that *tbx15* is responsive to CamK2b2levels, as injecting the CamKII expression construct alone led to *tbx15*upregulation. In addition, Tbx15 activity is sufficient to modulate any negative effects of increased CamKII activity, as the embryos resulting from injection of the CamKII expression construct do not have reduced numbers of pLL neuromasts unless increased CamKII activity is combined with *tbx15* knockdown. However, Camk2b2 is not solely responsible for *tbx15* activation, as we note *tbx15* upregulation in CamK2b2 morphants as well. Thus, CamK2b2 is sufficient but not necessary for *tbx15*activation, as it appears that Tbx5 is able to activate *tbx15* independent of Camk2b2, whether directly or via an alternate downstream target.

Data indicate that the *tbx5/tbx15* negative feedback loop is essential to a number of other pLL processes, as *tbx5* knockdown results in changes to multiple known primordium patterning markers; and indeed, all of our morphants demonstrate visible defects in primordium structure. Particularly, *tbx5* is implicated here in establishing differential expression of *cxcr4b* and *cxcr7b* and impacts overall expression levels for both. Furthermore, the *tbx5/tbx15* negative feedback loop appears at least partially responsible for expression of these chemokine receptors as we see that both *cxcr4b* and *cxcr7b* are downregulated in Tbx5 morphants and upregulated in Tbx15 morphants (Fig. 7). Interestingly, this feedback mechanism may explain the differences in severity between the Tbx5 and Tbx15 morphants. For example, *tbx5* knockdown may result in a more significant impact due to its direct role in activation of downstream targets. In contrast, the modulatory role of Tbx15 leads to a less severe phenotype, as transcriptional activation of downstream targets is still occurring, just not at the appropriate level. Of broader interest, we know that *tbx15* is expressed along with *tbx5* in the pectoral fin buds (Begemann et al., 2002), thus this feedback mechanism could be essential to the fin bud developmental process as well.

The actual mechanisms of Tbx5 activity remain unknown; however, our data along with previous publications provide clues as to the role or roles that Tbx5 occupies in pLL development. The impacts of Tbx5 on *cxcr4b* and *cxcr7b* expression provide an obvious mechanism for follow-up, as they along with their ligand, Sdf1a, are recognized for roles in primordium migration. In addition, Notch3 and Atoh1a activity in neuromast maturation and cell fate are important downstream effects that may be further investigated.

A study in chicken wing development showed that forelimb initiation and fate are dependent on tbx5 and that the impacts of tbx5 occur upstream of Wnt/FGF signaling (Takeuchi et al., 2003a). They further demonstrated loss of Fgf10 and Wnt2b following loss of Tbx5 (Takeuchi et al., 2003a). Thus, it is possible that *tbx5* is also important in setting up the Wnt/FGF signaling pathways that are essential to primordium patterning and migration in the developing pLL. Interestingly, we show *lef1* downregulation in Tbx5 morphants. Lef1 is a transcription factor that is activated by binding of stabilized ß-catenin in canonical Wnt/ß-catenin signaling. It has been implicated in several processes including mesoderm to epithelial transition, establishing Wnt signaling gradients and inhibition of Wnt signaling (Boras-Granic et al., 2006; Bazzi et al., 2007; Machon et al., 2007; Ling et al., 2009). It has also been suggested that Lef1 contributes to a Wnt signaling switch, turning off canonical pathways and turning on non-canonical pathways (Bazzi et al., 2007). Importantly, the Wnt/calcium pathway, which has the ability to activate CamKII, is associated with migration, and the Wnt/PCP pathway is essential in specification of hair cell orientation (Kuhl, 2004; Poggi et al., 2007; Jones and Chen, 2008; Samarzija et al., 2009). Thus, there are several avenues for investigating the significance of our finding that Tbx5 may upregulate *lef1*.

Though the roles of CamK2b2 in pLL development remain unclear, we do know that it is involved in negative regulation of *tbx5* via Tbx15. There is the potential for its involvement in signaling, migration, or both. Though we have shown that Tbx5 upregulates *camK2b2*, we don’t have evidence that it activates Camk2b2. Activation may be accomplished via other mechanisms. In this context, it is interesting that Tbx5 has been implicated as an upstream activator of Wnt signaling cascades. Perhaps Tbx5 upregulates *camK2b2* expression partly in preparation for use in Wnt/calcium signaling events. It would certainly be informative to conduct studies of CamK2b2 activation status after expression is upregulated or after expression of other Tbx5 downstream targets.

Further investigation is needed to determine the exact impacts of *tbx5* and *camK2b2* on pLL development; however, we have demonstrated here that Tbx15 has a modulatory role on *tbx5* and *camK2b2* expression. This and other recent publications highlight the need for negative feedback regulation to maintain proper balance of *tbx5* expression, which is essential for normal development (Camarata et al., 2010). We must elucidate the mechanism by which this feedback control occurs. It is possible that Tbx15 directly binds and inhibits Tbx5, or it could compete for binding T-box sites. It additionally remains to be determined if there are other targets of this feedback loop and if their expression levels also affect Tbx15.

## Acknowledgements

We thank A. Chitnis, P. Hernandez, S. Moody, T. Friedman, and J. Liang for helpful discussions. We also thank S. Tombes (Richmond, VA, USA) for reagents, S. Chandrasekharappa for equipment use, and S. Wincovitch for equipment and imaging assistance. This research was supported by the Intramural Research Program of the National Human Genome Research Institute, National Institutes of Health (SB).

## Supplemental Information

**Figure S1.**
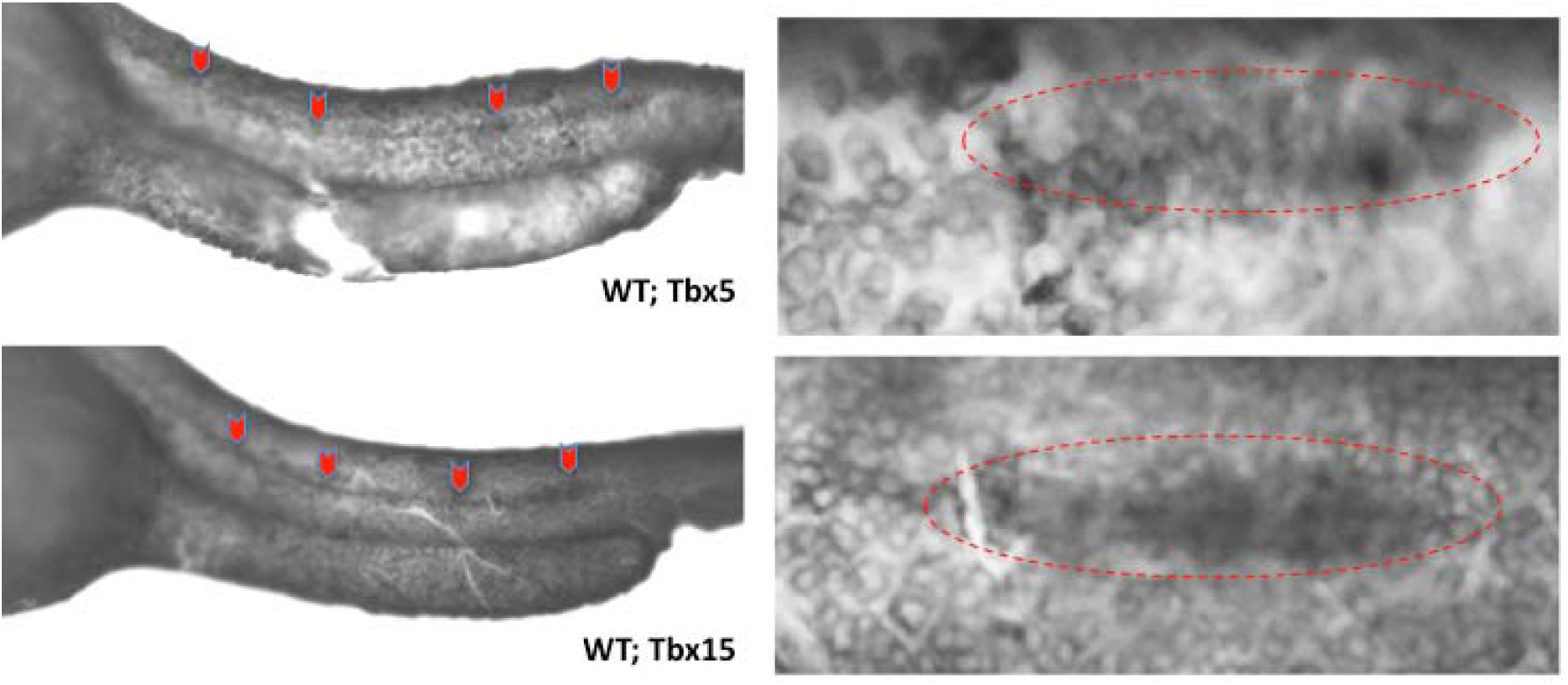
*tbx5* and *tbx15* are expressed in the migrating primordium and lateral line in wild-type embryos, and reduced expression is noted in morphants. Wild-type embryos were stained for Tbx5 and Tbx15 mRNA by in situ hybridization at 36 hpf. *tbx5* and *tbx15* expression are marked by enhanced staining along the pLL and in the pLL primordium.

**Figure S2.**
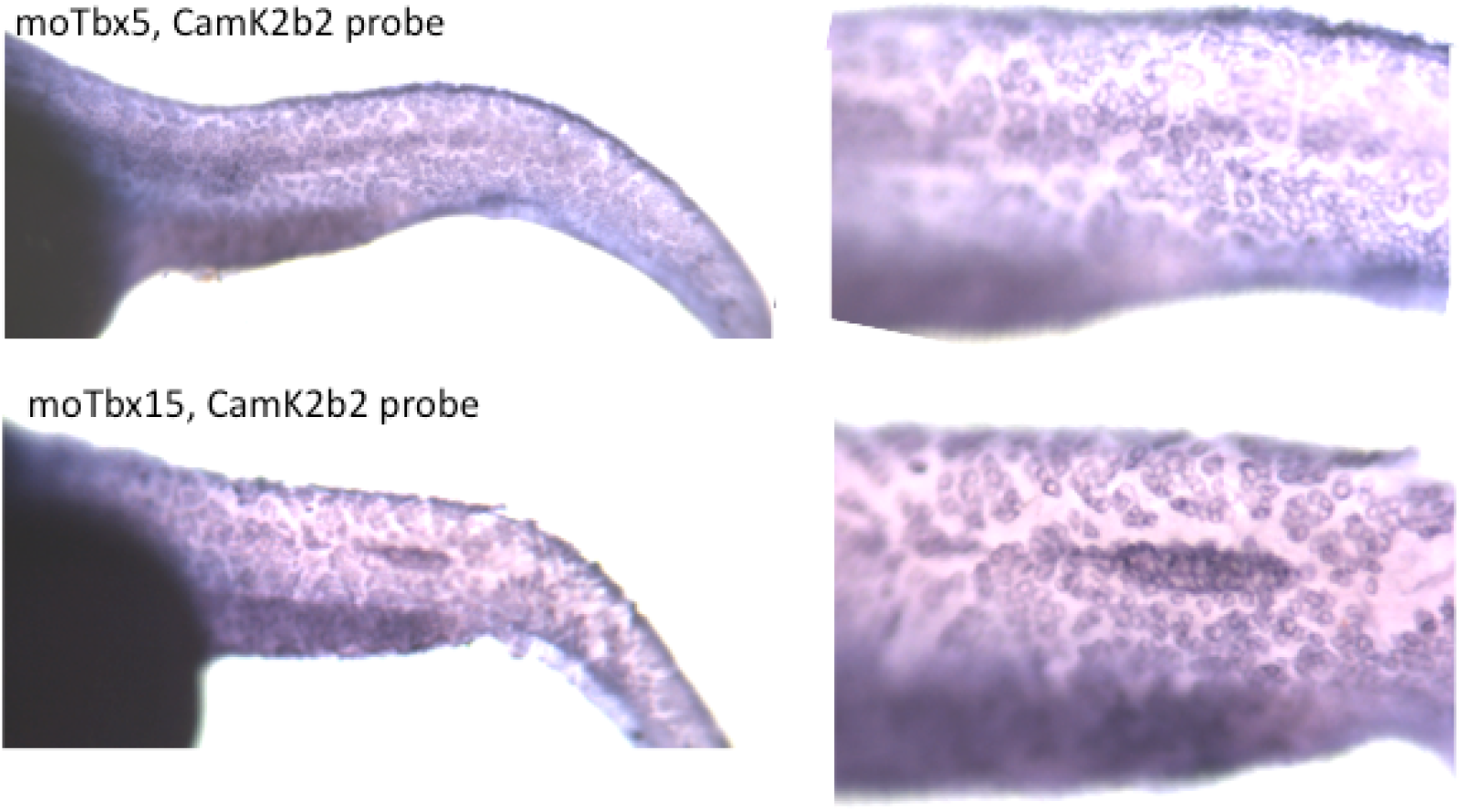
*camK2b2* expression is lost in Tbx5 morphants and is retained in Tbx15 morphants. *In situ* hybridization was done in Tbx5 morphants (top two panels) and Tbx15 morphants (bottom two panels) with digoxigenin-labeled RNA probes against Camk2b mRNA. Detectable levels of CamK2b2 mRNA are not found in Tbx5 morphants but are present in Tbx15 morphants.

**Figure S3.**
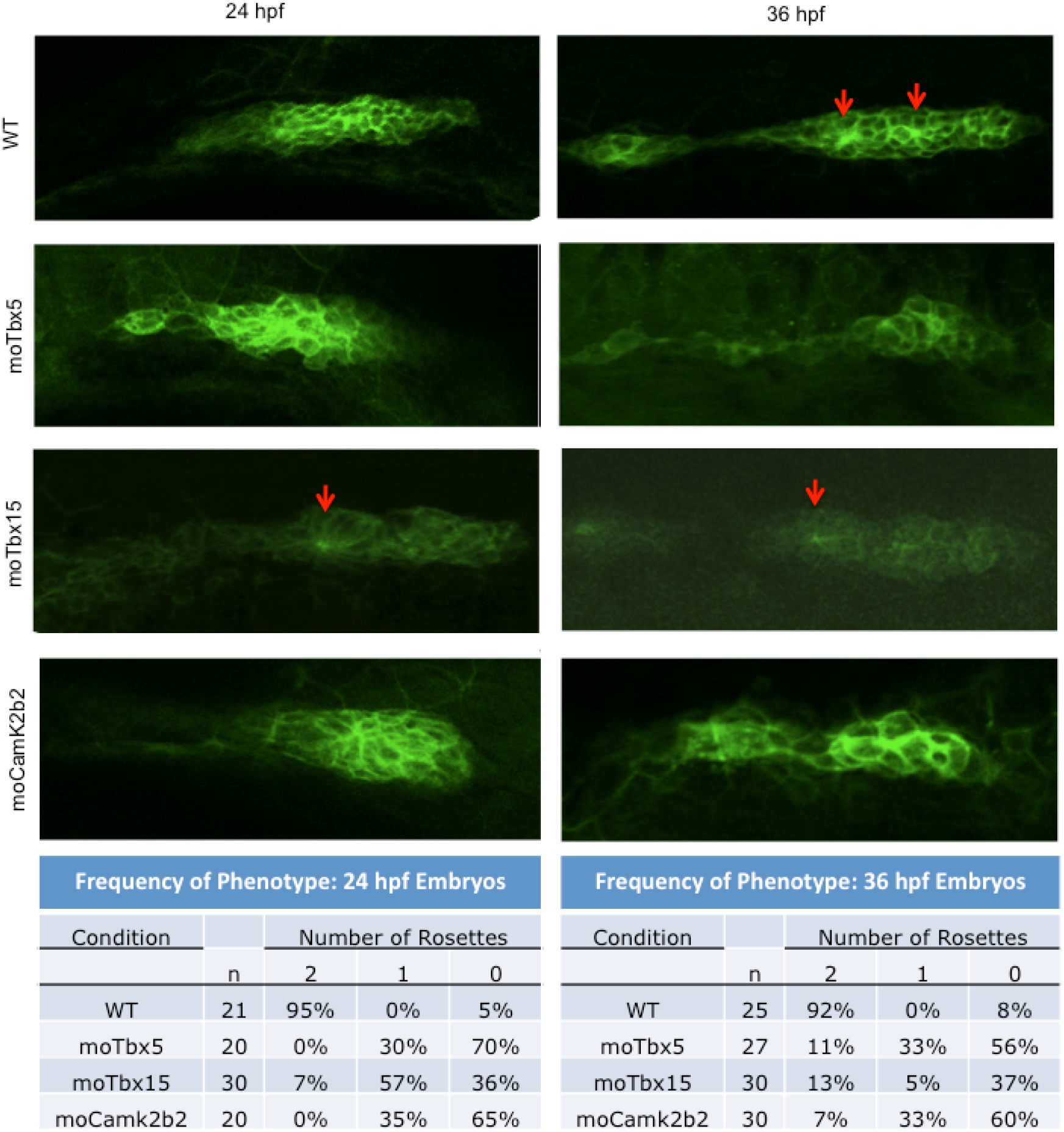
Primordium form and patterning are altered in Tbx5, Tbx15, and CamK2b2 morphants. WT, Tbx5, Tbx15, and CamK2b2 Tg(−8.0cldnb:lynEGFP) morphants were stained at 24 hpf and 36 hpf with a FITC-conjugated anti-GFP antibody and used for confocal imaging. The top two panels are migrating primordia from WT embryos. In the 36 hpf image, focal points indicating the centers of rosette structures within the primordium are noted (arrows). Overall form is impacted in Tbx5 morphants at 24 hpf and is lost by 36 hpf as are focal points indicative of rosettes. Sample sizes and phenotypes are summarized for each condition and embryo age.

**Figure S4.**
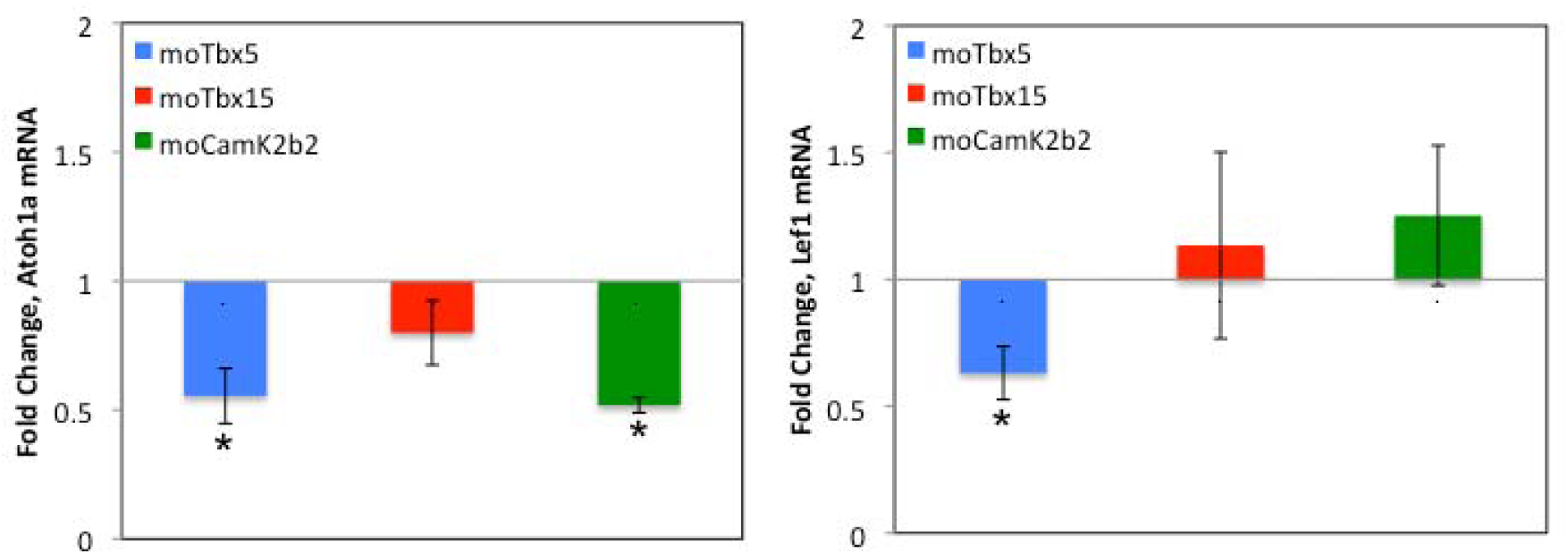
Markers of primordium patterning have altered expression levels in Tbx5, Tbx15 and Camk2b2 morphants. qRT-PCR analysis was conducted on WT, Tbx5, Tbx15 and Camk2b2 morphants at 36 hpf for Atoh1a and Lef1 mRNA. Expression of *atoh1a* is significantly reduced in both Tbx5 (p(tbx5) < 0.05) and Camk2b2 morphants (p(camk2b2) < 0.05). *atoh1a* expression is not significantly changed in Tbx15 morphants. Expression of *lef1* is significantly reduced in Tbx5 (p(tbx5) < 0.05) morphants. *lef1* expression is not significantly changed in Tbx15 or Camk2b2 morphants.

